# FKBP12-BOUND CALCINEURIN CONTROL TMEM16A ACTIVITY

**DOI:** 10.1101/2024.06.27.601078

**Authors:** María Luisa Guzmán-Hernández, Miriam Huerta, Ana E. López-Romero, Rita Morán-Zendejas, Abigail Betanzos, Patricia Pérez-Cornejo, Jorge Arreola

## Abstract

The Ca^2+^-dependent Cl^-^ channel TMEM16A activity is vital for mammals. Critical functions such as blood pressure, pain, fluid and electrolyte secretion, peristalsis, and electrical activity depend on TMEM16A Cl^-^ fluxes. Ciclosporin A (CsA) and FK506, two immunosuppressants that inhibit calcineurin (CaN), down-regulate TMEM16A function. However, the underlying mechanism remains undetermined. CsA binds to cyclophilin 1, whereas FK506 binds to FKBP12, and both complexes subsequently inhibit CaN, a phosphatase that maintains TMEM16A function. Here, we show that TMEM16A-EGFP and mRFP-FKBP12 colocalize in HEK-AD293 cells stimulated with the Ca^2+^ ionophore ionomycin, FKBP12 co-immunoprecipitated with TMEM16A expressed in CaN-depleted HEK-AD293 cells, ionomycin favoured TMEM16A and FKBP12 interaction in a bimolecular fluorescence complementation assay in live cells, and TMEM16A currents recorded from CaN-depleted HEK-AD293 cells were insensitive to CsA and FK506. These data support the idea that FKBP12 works as an auxiliary protein of TMEM16A, thus enabling the interaction of CaN under physiological conditions. This heteromerization is essential to sustain TMEM16A activity. On the contrary, disrupting the Ca^2+^-stimulated CaN-FKBP12-TMEM16A heterotrimer or inhibiting CaN with CsA or FK506 does not preclude TMEM16A activation but decreases activity.

## Introduction

TMEM16A is a Cl^-^ channel activated by increments in the cytosolic free Ca^2+^ concentration induced by activation of G protein-coupled receptors or activation of Ca^2+^ channels (Caputo et al., 2008; Schroeder et al., 2008; Yang et al., 2008). The Cl^-^ fluxes via TMEM16A are essential to maintain and carry on vital physiological functions (Pedemonte and Galietta, 2014; Arreola et al., 2022; Hartzell et al., 2009) including blood pressure, transduction of painful stimuli (Cho et al., 2012; Lee et al., 2014), fluid and electrolyte secretion (Romanenko et al., 2010; Huang et al., 2012; Namkung et al., 2011), and hormone secretion (Crutzen et al., 2016; Xu et al., 2014). On the other hand, when TMEM16A is overexpressed or mutated, harmful diseases, such as hypertension (Askew Page et al., 2019; Forrest et al., 2012), cancer and tracheomalacia (Crottès and Jan, 2019; Duvvuri et al., 2012; Rock et al., 2008) are developed. The Ca^2+^-dependent function of TMEM16A has been extensively studied at cellular, molecular, and structural levels (Arreola et al., 2024). It is now well established that Ca^2+^ triggers TMEM16A activity by interacting with at least three binding sites. Two of these sites have been molecularly and structurally identified (Paulino et al., 2017; Yu et al., 2012; Tien et al., 2014; Dang et al., 2017). Residues from helices 6, 7, and 8 shape the orthosteric site, while the third site is located at the dimer’s interface and formed by residues from helices 2 and 10 (Le and Yang, 2020). When Ca^2+^ binds to the orthosteric site, a chain of structural rearrangements triggers pore opening and Cl^-^ conduction. (Paulino et al., 2017; Lam et al., 2021; Lam and Dutzler, 2021).

The cellular and molecular mechanisms that control TMEM16A function are less understood. It has been proposed that the phosphorylation/dephosphorylation balance of TMEM16A is critical for its activity (Leblanc et al., 2005). CaMKII-mediated phosphorylation diminishes TMEM16A activity, whereas phosphatase-dependent dephosphorylation augments activity (Angermann et al., 2006; Ayon et al., 2009, 2019; Wang and Kotlikoff, 1997). It has been proposed that calcineurin (CaN) regulates TMEM16A by a dephosphorylation mechanism. In agreement, the Ca^2+^-activated Cl^-^ currents in coronary arteries were decreased by CaN inhibitors, such as CsA, whereas in pulmonary arteries increased by the constitutively active CaN (Greenwood et al., 2004a; Ledoux et al., 2003). A common feature of TMEM16A, CaMKII, and CaN activation processes is their Ca^2+^ dependence. These proteins are inactive at resting cellular Ca^2+^ levels, yet their activity rises gradually as intracellular Ca^2+^ increases after a physiological stimulus (REF). Previously, we reported that under high intracellular Ca^2+^, TMEM16A interacts with CaN and FKBP12. In addition, ciclosporin A (CsA) and FK506 diminished but did not abolish TMEM16A activity (Sánchez-Solano et al., 2020). We hypothesize that under physiological conditions, an increase in intracellular Ca^2+^ induces the formation of the hetero-trimeric complex TMEM16A-FKBP12-CaN and that this complex is essential to sustain TMEM16A activity. In addition, FK506 could inhibit Ca^2+^-dependent Cl^-^ secretion in airway epithelium via a CaN-independent mechanism (Kanoh et al., 2001). In this work, we determined the interaction and functional alterations of TMEM16A expressed in HEK-AD293 cells transfected with a shRNA directed against CaN. Our data show that in this cellular background, TMEM16A interacted with FKBP12 and that CsA and FK506 did not further diminish TMEM16A activity, indicating that CaN mediates their actions. We also demonstrate that FKBP12 interacts directly with TMEM16A in intact cells and membrane patches attached to coverslips. Our work reveals a regulatory mechanism of TMEM16A activity whereby the CaN-FKBP12 pair interacts to up-regulate the chloride current.

## Materials and methods

### Cell culture and transfection

HEK293 and HEK-AD293 cells (ATCC, Manassas, VA, USA) were cultured in Dulbecco’s modified Eagle’s medium (DMEM) supplemented with 10% (v/v) fetal bovine serum (FBS; both from GIBCO, Carlsbad, CA, USA), and 0.1% penicillin-streptomycin (Sigma-Aldrich). Stably transfected HEK-AD293 cells were maintained in DMEM supplemented with 0.6 μg/ml puromycin (Sigma-Aldrich), 10% FBS and 0.1% penicillin-streptomycin. All cell cultures were incubated at 37 °C in a humidified atmosphere with 5% CO_2_. Transient transfections using Lipofectamine 2000 reagent and OPTI-MEM (both from Invitrogen) were carried out according to the manufacturer’s instructions.

### Stably transfected HEK-AD293 cells

HEK-AD293 cells were seeded onto 35-mm diameter dishes and cultured until they reached 40-50% confluency. To obtain a stable cell line with reduced CaN expression, we transfected the cells with a small hairpin RNA (shRNA) vector against human calcineurin A (PPP3CA), which was cloned in a lentiviral plasmid carrying a green fluorescence protein (GFP) tag (cat#: TL310220, OriGene Technologies, Inc.). Cellular dilutions followed transfections using Lipofectamine 2000 for three days to facilitate cell isolation. Puromycin was added to select cells expressing shRNA against CaN or scrambled shRNA (control cells). Isolated colonies were subsequently recovered using cloning rings and seeded into individual wells. Cell fluorescence was monitored over a six-week culture period, and silencing of CaN expression was assessed by Western blotting. The resulting stable lines were designated as CaN KD and Ctrl KD.

### Total protein extraction from stably transfected HEK-AD293

The CaN KD and Ctrl KD cells were rinsed three times with phosphate-buffered saline (PBS) and then stimulated with 1 μM Ionomycin (I9657, Sigma-Aldrich) for 30 s in the presence of 1X PBS containing 0.5 mM CaCl_2_. Following stimulation, cells were placed on ice and lysed with 50 μL of buffer containing (in mM) 150 NaCl, 10 HEPES, 1 EGTA, and 0.1 MgCl_2_, pH 7.4, supplemented with 0.5% Triton X-100 and protease inhibitors (Complete Ultra, Roche). Lysates were incubated for 30 minutes at 4 °C, followed by centrifugation at 14,800 rpm for 15 min at 4 °C. The supernatant, containing the total protein lysate, was recovered and stored at -80°C. Protein concentration was determined using the Bradford method.

### Immunoprecipitation and Western blot assays

The interaction between TMEM16A and FKBP12 was assessed in CaN KD and Ctrl KD cell lines seeded onto 10-cm diameter dishes. Cell lines were transfected with 1 μg of pEGFP-N1 plasmid containing the cDNA encoding mouse TMEM16A (mTMEM16A, ac variant) tagged with 3XFlag and 24 h after transfection protein complexes were immunoprecipitated (Sánchez-Solano et al., 2020). Total protein (1000 μg) was incubated for 3 h at 4 °C with 1 μl of mouse anti-Flag M2 antibody or 3X Flag peptide (200 mg/ml; Sigma-Aldrich) on a rocking platform. Then, 30 μl of protein A/G plus-agarose beads (Santa Cruz Biotechnology) were added, and the cell lysates were incubated for 12 h at 4 °C. The beads were washed five times with 1 ml of ice-cold lysis buffer. Immunocomplexes were eluted using 35 μl Laemmli buffer 2X and boiled at 95 °C for 5 min. 35 μl of immunoprecipitated proteins were recovered and loaded into the gel. Proteins were separated by SDS-PAGE and transferred to PVDF membranes using a semi-dry transfer cell (Bio-Rad, Hercules, CA, USA). Membranes were blocked with 5% non-fat dry milk in TBS-Tween for 1 h at room temperature, and proteins were detected by Western blot using monoclonal anti-Flag M2 antibody produced in mouse (1:5000 dilution; Sigma-Aldrich), rabbit anti-FKBP12 antibody (1:1000 dilution; Abcam) and rabbit anti-calcineurin A antibody (1:5000 dilution; Abcam). GAPDH was used as an internal loading control. A mouse anti-GAPDH antibody (1:20,000 dilution; Santa Cruz Biotechnology) was utilized to detect GAPDH expression in total lysates. Goat anti-mouse (1:5000 dilution; Millipore) and mouse anti-rabbit (1:5000 dilution; Santa Cruz Biotechnologies) were employed as secondary antibodies. Immunoblots were visualized by chemiluminescence using the ChemiDoc XRS+ system and Image Lab Software (Bio-Rad).

### Co-localization assays

HEK-AD293 cells were seeded onto 1 cm2 glass coverslips coated with poly-L-Lysine and placed in a 24-well plate (20,000 cells/well) to be co-transfected with 1.5 μg EGFP-TMEM16A plasmid (kindly donated by Dr. Utaek Oh) and 0.5 μg mRFP-FKBP12 plasmid (kindly donated by Dr. Tamás Balla). 48 h post-transfection, cells were treated with either PBS, PBS+Ca^2+^ (0.5 mM CaCl_2_), or 1 μM Ionomycin for 30 s or 5 min. Next, cells were fixed with 4% paraformaldehyde for 10 min in cold conditions and extensively washed. Coverslips were mounted using Fluoromount Aqueous Mounting Medium (Sigma-Aldrich) and later examined using a Carl Zeiss LMS 700 confocal microscope in Z-stack to obtain laser sections of 0.5 μm thickness. Excitation/emission wavelengths of 410/520 and 603/700 nm were used to detect EGFP and mRFP, respectively. Images were processed with ZEN 3.8 Light Edition software (Carl Zeiss Microscopy GmbH).

### Bimolecular Fluorescence Complementation (BiFC)

The BiFC assay was conducted as described by Kodama et al (Kodama and Hu, 2012). The N-terminus of FKBP12 was fused to amino acids 1– 154 of the fluorescent protein Venus (VN-FKBP12), which includes the I152L point mutation to reduce self-assembly. Likewise, the C-terminus of TMEM16A was fused to amino acids 155–239 of Venus (TMEM16A-VC). HEK-AD293 cells seeded onto poly-L-Lysine-coated coverslips in 24-well plates (15,000 cells/well) were first transfected with 250 ng TMEM16A-VC and, after 12h, co-transfected with 250 ng VN-FKBP12 and 150 ng mCherry empty vector. We used mCherry to find successful transfected cells and mRFP-FKBP12 as a negative control. Following a 24-hour post-transfection period, cells were treated with PBS or PBS containing 0.5 mM Ca^2+^ and 1 μM ionomycin for 50 min at 37 °C. To analyze complementation between TMEM16A-VC and VN-FKBP12 in living cells, a coverslip was placed into a chamber, and the cells were visualized using the Olympus IX73 fluorescence microscope equipped with a Hamamatsu Digital Camera C13440 ORCA-Flash 4.0 and a xenon arc lamp Lambda LS. BiFC signal was collected at 507-543 nm from cells excited at 480-500 nm. mRFP fluorescence was collected using the Texas-Red filter (excitation 540-580 nm and emission 592-667 nm). Images were acquired using the CellSens software and analyzed using Image J.

### Electrophysiological recordings

HEK293 cells were seeded at low density onto coverslips pre-treated with poly-L-lysine (Sigma-Aldrich) and recorded using the whole-cell configuration of the patch clamp technique. Cells were transfected with 800 ng of mouse TMEM16A (ac) cloned into a mCherry vector (TMEM16A-mCherry, kindly donated by Dr. C. Hartzell) and recorded 18 h post-transfection. For knock-down experiments, cells were transiently transfected with 1 μg of shRNA against CaN or scrambled shRNA 72 h prior to transfection with the TMEM16A-mCherry construct.

The extracellular solution contained (in mM) 139 TEA-Cl, 20 HEPES, 0.5 CaCl_2_, and 110 D-mannitol. The pH was adjusted to 7.3 with TEA-OH. The solution’s tonicity was 380–400 mOsm/kg, measured by the vapour pressure point method with a VAPRO Osmometer (Wescor). The intracellular solution contained (in mM): 30 TEA-Cl, 25 EGTA-TEA, 5.24 CaCl_2_, 50 HEPES and 85 D-Mannitol and a final 0.2 μM free Ca^2+^ concentration, which was calculated using the Maxchelator online calculator (https://somapp.ucdmc.ucdavis.edu/pharmacology/bers/maxchelator/). The pH was adjusted to 7.3 with TEA-OH and the tonicity to 290–300 mOsm/kg. Cyclosporin A (CsA) stock solution was prepared at 10 mM. Cells were pre-treated for 1 h with 20 μM CsA and recorded in the presence of 20 μM CsA in the pipette solution. All reagents were purchased from Sigma-Aldrich.

Cells displaying mCherry and GFP fluorescence were positive for channel and shRNA expression and were selected using an epifluorescence inverted Nikon microscope equipped with a UV lamp. Borosilicate glass microelectrodes were fabricated using the P-97 microelectrode puller (Sutter Instruments). Currents were recorded with an Axopatch 200B amplifier at room temperature (21-23°C) by applying from a holding potential of -20 mV a voltage step protocol that consisted of 500 ms voltage steps that varied from -100 mV to +160 mV in 20 mV increments, and a repolarizing voltage of -100 mV that lasted 250 ms. The recording chamber was grounded using a 3 M KCl agar bridge connected to an Ag/AgCl reference electrode, and the bath solutions were exchanged using a gravity perfusion system. Data were filtered at 5 kHz and digitized at 10 kHz using an Axon Digidata 1550 and the pClamp10 software (Molecular Devices).

### Analysis

For co-localization experiments, the degree of TMEM16A and FKBP12 signal overlap was inferred by calculating the Manders Pearson’s coefficient from the middle image of the Z-stack in both channels employing the ImageJ 1.54d software and JACoP (Just Another Colocalization Plugin) plugin (https://imagej.net/ij/plugins/track/jacop2). In this analysis, the coefficient values can vary from -1 to 1, with -1 meaning a negative correlation, 0 no correlation, and 1 complete positive correlation. In three independent experiments, four equal-sized regions, three at the membrane and one at the cytosol, were analyzed for each cell taken from a collection of at least 15 cells. Fluorescent profiles were obtained using the ZEN software. Statistical differences between the Pearson’s coefficients of the groups were evaluated using a one-way ANOVA, followed by Tukey’s multiple comparison test.

The bimolecular fluorescence complementation (BiFC) assay was analyzed as follows. Cells from each experiment were quantified using ImageJ. Cells from three experiments were analyzed, and BiFC measurements were presented as the whole-cell mean fluorescence values. Statistical differences were evaluated using a one-way ANOVA test.

CaN and GAPDH Western blot band intensities were detected using the Image Lab Software (BioRad). CaN band intensities were normalized using the band intensities of the GAPDH, the loading control, as follows. We divided the GAPDH signal intensity in the control condition by the GAPDH band intensities in the control and experimental conditions to get the normalization factor for each condition. Then, CaN band intensities were scaled by the corresponding normalization factor. For co-immunoprecipitation experiments, only the signal intensity observed in the blot is reported.

Current versus voltage relationships were constructed using the average current density (pA/pF). The magnitude of whole-cell current was measured at the end of each voltage step, normalized to the cell’s capacitance, pooled at each voltage, and then averaged. The current density is presented as mean ± SEM.

Origin software was used to plot and perform the statistical analysis. *n* represents the number of cells recorded (in patch clamp) or the number of independent experiments.

## Results

### Calcineurin is dispensable for FKBP12 binding to TMEM16A

Previously, we reported that TMEM16A could form a ternary complex with FKBP12 and CaN in a Ca^2+^-dependent manner (Sánchez-Solano et al., 2020). To further elucidate the role of CaN in the interaction between TMEM16A and FKBP12, we generated a CaN knockdown cell line (CaN KD) and assessed complex formation by co-immunoprecipitation. HEK-AD293 cells transfected with short hairpin RNA (shRNA) targeting CaN were subjected to puromycin selection and monitored by looking at the fluorescence of the GFP-tagged plasmid that contained the shRNA vector. The silencing of CaN was confirmed by Western blot analysis. As depicted in Fig. 1A, CaN expression was significantly reduced, while cells transfected with a control scramble shRNA, which does not target any specific gene, showed a robust CaN expression. The band intensity corresponding to CaN decreased 11-fold in the total lysate of CaN KD cells (Fig. 1B). Next, we investigated whether, in the absence of CaN, the interaction between TMEM16A and FKBP12 persisted. Both control KD and CaN KD cells were transfected with TMEM16A-3XFlag (ac variant) cDNA. After 24 h, the culture media was replaced with a solution containing 0.5 mM Ca^2+^, and the cells were exposed to 1 μM ionomycin for 30 s; immediately after, the cells were lysed for protein isolation. As depicted in Fig. 1C, TMEM16A coimmunoprecipitated with FKBP12 irrespective of the presence (left lane, control cells) or absence of CaN (middle lane, CaN KD cells). As a control, in some experiments, the 3X-Flag peptide was added to compete for the anti-FLAG antibody, decreasing the amount of TMEM16A-3XFlag bound to the agarose beads (right lane). After quantification, the FKBP12 signal intensity from CaN KD cells was not different from that of immunoprecipitation experiments with control cells or the 3X-Flag peptide (Fig. 1D). However, using the 3X-Flag peptide diminished FKBP12 recognition by 45%. Figure 1C also shows the expression of TMEM16A and FKBP12 in the protein lysates used for CoIP experiments. The expression of these proteins remained unaffected by CaN shRNA. These findings suggest that CaN contributes to the trimeric complex but is not essential for the interaction between FKBP12 and TMEM16A.

**FIGURE 1.**
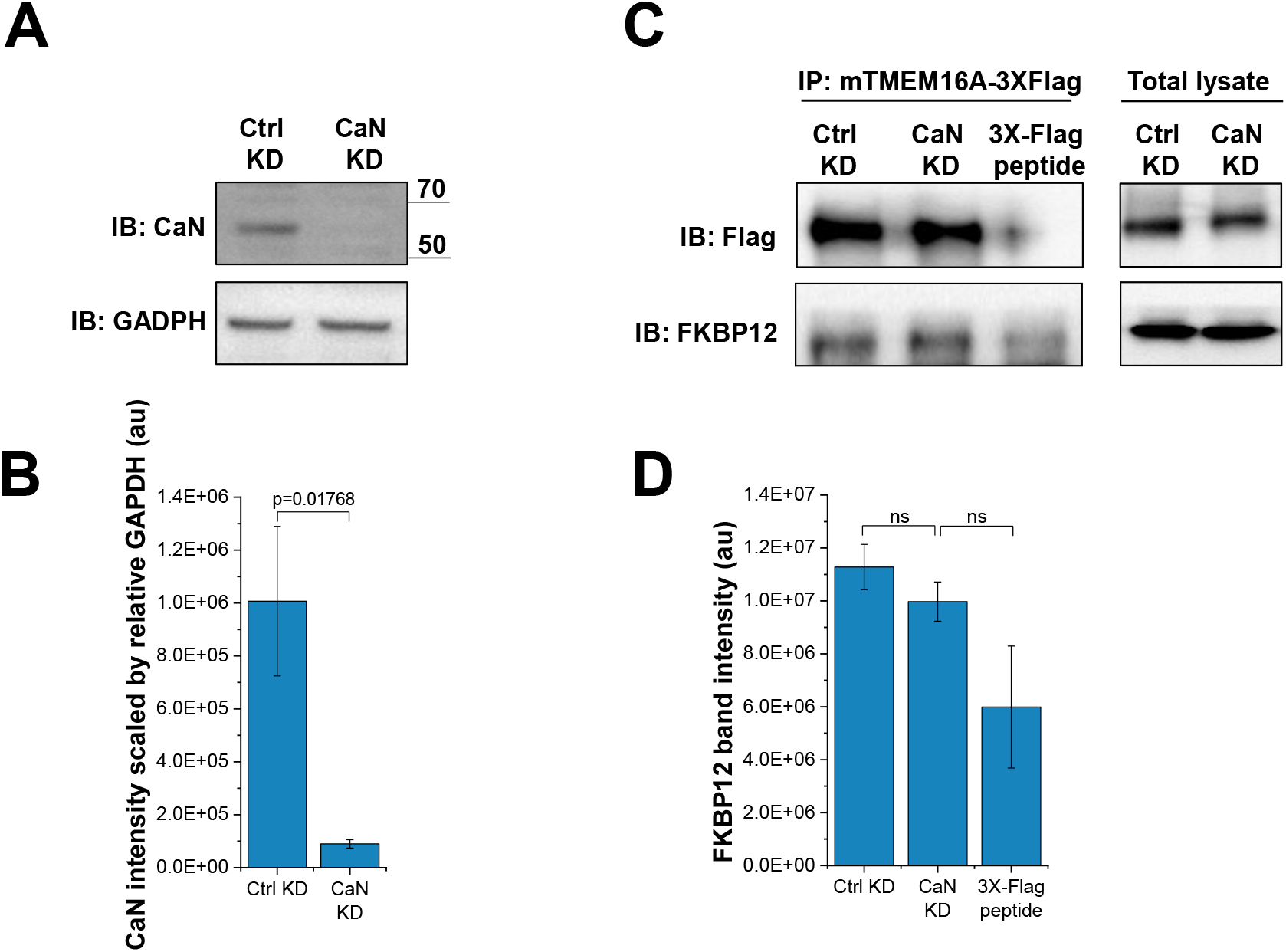
CaN is dispensable in the interaction between TMEM16A and FKBP12. **A)** Knockdown of CaN in stably transfected HEK-AD293 cells (CaN KD). Protein samples were probed with anti-CaN antibodies, revealing a diminished CaN KD cell line expression compared to the Ctrl KD. A representative blot from three independent experiments is shown. **B)** Quantification of CaN band intensities scaled by the relative GAPDH. Mean values ± SEM are presented (n=4). **C)** TMEM16A-3XFlag was immunoprecipitated, and FKBP12 was detected in both CaN KD and Ctrl KD cell lines. An interaction between TMEM16A and FKBP12 is observed in the absence of CaN when intracellular Ca^2+^ levels are increased with 1 μM ionomycin, but this interaction is weaker in the presence of the 3XFlag peptide. A representative blot from three independent experiments is provided. Protein expression in total lysate for TMEM16A and FKBP12 is shown. **D)** Quantification of FKBP12 band intensities. Mean ± SEM, (n=3).

### TMEM16A co-localizes with FKBP12

Given that CaN is not necessary for the TMEM16A-FKBP12 interaction, we explored the association between TMEM16A and FKBP12 *in vivo* by co-localizing both proteins in HEK-AD293 cells. TMEM16A tagged with EGFP and FKBP12 tagged with mRFP were co-transfected. After 48 h, the cells were incubated with either PBS or PBS containing 0.5 mM Ca2+ and then stimulated with 1 μM ionomycin for 30 s and 5 min to increase the intracellular Ca^2+^ concentration and promote TMEM16A activation. As shown in Fig. 2A, the TMEM16A-EGFP signal localized to membrane regions (left lane), whereas mRFP-FKPB12 was distributed throughout the cell (center lane). Combining the signals from both channels (right lane) revealed a yellow signal, mostly on cells treated with ionomycin. This signal became stronger after 5 min of incubation (fourth row). These observations are summarized as the fluorescence profile of each representative image (Fig. 2B).

**FIGURE 2.**
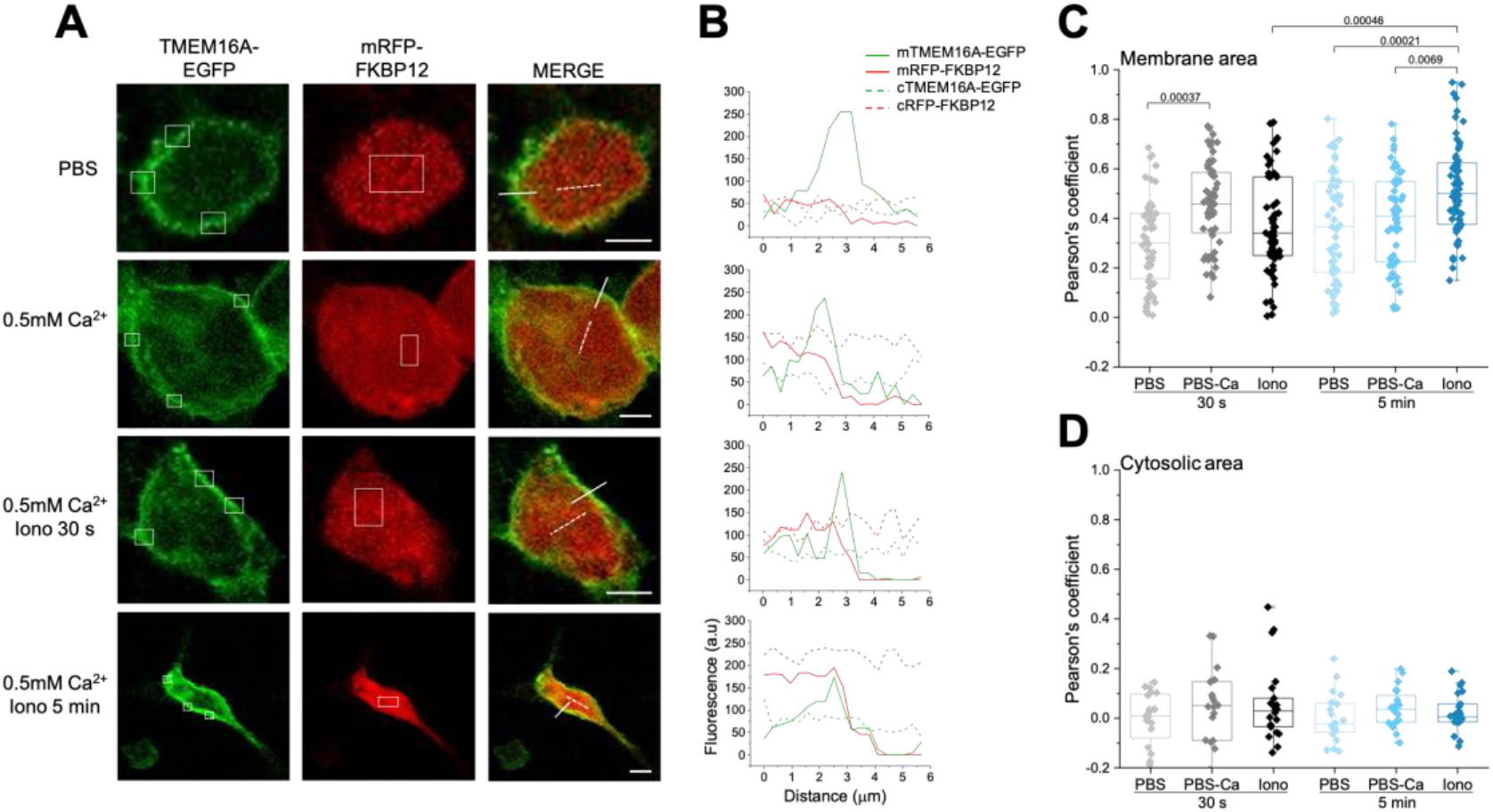
TMEM16A and FKBP12 co-localized at the cell membrane after ionomycin exposure. **A)** Representative confocal images of HEK293AD cells co-expressing TMEM16A-EGFP and mRFP-FKBP12 proteins after treatments with PBS (first row, negative control), PBS + 0.5 mM Ca2+ (second row), and 1 μM Ionomycin in 0.5 mM Ca^2+^ during 30 s (third row) and 5 min (fourth row). Rectangles of equal area (left panels) were placed on three different points at the membrane and one more was placed in the cytoplasm (middle panels) to analyze the fluorescence in each cell. Scale bar = 5 μm. **B)** Representative fluorescence profiles for TMEM16A (green line) and FKBP12 (red line) proteins in co-transfected cells. Continuous lines represent the signal from the membrane (m) and dashed lines the signal from the cytosol (c). Fluorescence profiles were constructed measuring the fluorescence along the continuous (membrane) and dashed (cytosol) lines shown on the merge column. **C)** Pearson’s coefficient was calculated for each experimental condition and these values were used to infer TMEM16A-FKBP12 co-localization. The analysis was performed in 15 cells corresponding to 45 membrane (panel C) and 15 cytosolic regions (panel D). The p values for the statistical comparisons are shown above the boxes.

To quantify the co-localization of TMEM16A and FKBP12, we calculated Pearson’s correlation coefficient, as indicated in the Methods section. Since we noticed that the TMEM16A signal was not homogeneous, we analyzed three equal areas in different cell membrane regions to avoid a potential underestimation of Pearson’s coefficient values. There seems to be some co-localization in PBS-treated cells, but we observed a slight rise in Pearson’s values after adding Ca^2+^ to the external solution and waiting 30 s. The treatment with ionomycin did not affect the initial co-localization; however, we observed two different populations, suggesting that not all cells and regions were responsive to Ionomycin. Interestingly, Ionomycin-induced co-localization became evident and statistically significant after a 5 min treatment (Fig. 2C). Pearson’s coefficient was near zero in cytosolic regions, indicating no interaction of the proteins in this region (Fig. 2D). Taken together, our findings suggest that the membrane-delimited TMEM16A-FKBP12 interaction intensifies following an intracellular rise in Ca^2+^.

### Intracellular Ca^2+^ enhances the TMEM16A-FKBP12 interaction in live cells

Our co-localization experiments suggest that when intracellular Ca^2+^ increases, FKBP12 is redistributed closer to the membrane. Thus, we carried out the bimolecular fluorescence complementation (BiFC) assay to determine a direct interaction between these proteins in living cells. These experiments were performed in the presence of 1µM ionomycin and 0.5 mM Ca^2+^. To this end, we fused amino acids 1– 154 and 155–239 of the Venus protein to the N- and C-terminus of FKBP12 and TMEM16A, respectively. If FKBP12 relocates to the membrane and binds TMEM16A, structural complementation between Venus’s N- and C-terminal fragments would occur. As a result, we could record Venus’s fluorescence at 507-543 nm. As depicted in Fig. 3A, the co-expression of VN-FKBP12 and TMEM16A-VC in HEK-AD293 cells resulted in a fluorescence signal generated by Venus’s complementation (bottom left lane). This indicated that VN-FKBP12 had relocated to the plasma membrane and could interact with TMEM16A-VC. Despite the slow maturation of the Venus protein after complementation (_∼_1h), a stable and functional complex between FKBP12-TMEM16A was observed. In contrast, when we transfected mRFP-FKBP12, a protein lacking the N-terminus half of Venus, along with TMEM16A-VC (middle), the total fluorescence recorded was significantly reduced. Results from an additional control competition assay, in which mRFP-FKBP12 was put to compete with VN-FKBP12 for interaction with TMEM16A-VC, show a significant decrease in the BiFC signal (bottom right). The transfection of HEK cells with a vector carrying either the N or V terminus of Venus alone did not produce a BiFC signal. This shows that a BiFC signal is observed only after TMEM16A-VC and VN-FKBP12 interact. The statistical analysis of the images indicated that the association between FKBP12 and TMEM16A is significantly different from the control experiments (Fig. 3B). Interestingly, we found a significant basal association between FKBP12 and TMEM16A compared with control conditions in which the cells were not treated with ionomycin (Fig. 3C and 3D). The Venus BiFC data show that the FKBP12-TMEM16A interaction occurs in living cells.

**FIGURE 3.**
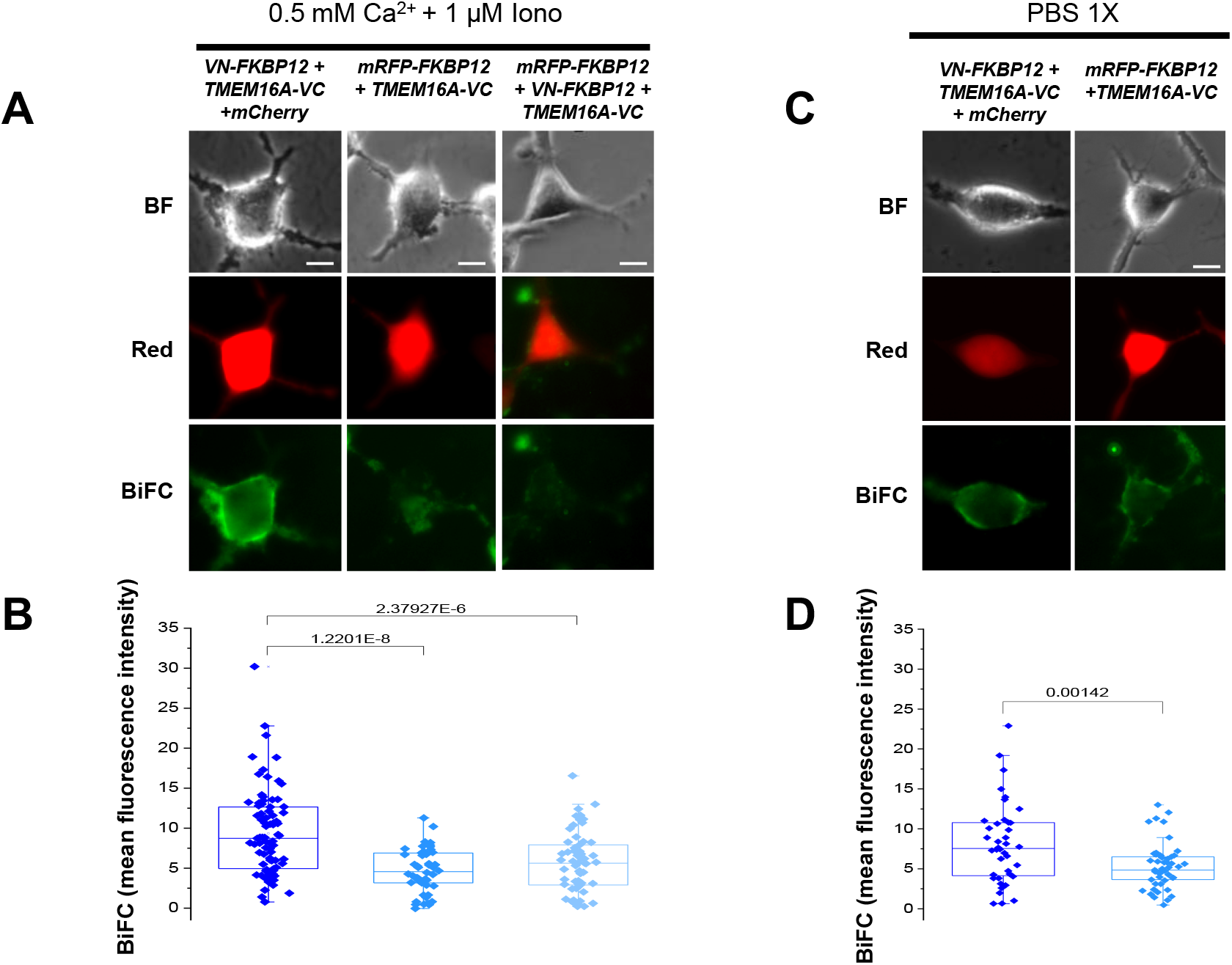
FKBP12 and TMEM16A interact in live cells. **A)** Representative images of HEK-AD293 cells transfected with TMEM16-VC + VN-FKBP12 plus mCherry empty vector (left), TMEM16A-VC and mRFP-FKBP12 (middle), or TMEM16A-VC + VN-FKBP12, and mRFP-FKBP12 (right). Images in red are from mCherry or mRFP-FKBP12 fluorescence. Images in green show the BiFC signal indicative of Venus complementation. All images were taken in the presence of 1µM Ionomycin and 0.5mM Ca^2+^ in the extracellular solution. **B)** Mean BiFC fluorescence intensities measured from images like those in A. Statistical analysis revealed that BiFC signal was significantly higher in cells transfected with TMEM16-VC + VN-FKBP12, and mCherry than in control conditions (without VN-FKBP12: p-value = 1.22E-8, n = 83; competition: p-value = 2.37E-6, n = 60). **C)** Representative fluorescence images taken from two cells transfected with TMEM16-VC + VN-FKBP12 plus mCherry empty vector (left lane) or TMEM16A-VC and mRFP-FKBP12 (right lane). These cells were not exposed to ionomycin. **D)** Mean BiFC fluorescence intensities measured from images like those in C. Statistical analysis revealed that BiFC signal was significantly higher in cells transfected with TMEM16-VC + VN-FKBP12 plus mCherry than control condition without VN-FKBP12 (p-value = 0.00142, n = 50). This result show that FKBP12 and TMEM16A form a stable basal complex in living cells. Bar size = 10 μm. Statistical differences between data sets were evaluated by a one-way ANOVA statistical test.

### Calcineurin regulates TMEM16A activity

CaN interacts with FKBP12 and TMEM16A when the intracellular Ca^2+^ increases (Sánchez-Solano et al., 2020), furthermore CaN regulates TMEM16A activity (Greenwood et al., 2004b). The association between FKBP12 and TMEM16A is not affected by the absence of CaN, but is CaN still required to regulate TMEM16A activity? To answer this question, we performed patch-clamp experiments in HEK293cells expressing TMEM16A-mCherry that were untransfected (wild type), transfected with shRNA against CaN (CaN KD), or transfected with scrambled shRNA (scrambled). Moreover, we tested two CaN inhibitors: CsA, which forms a complex with cyclophilins and inhibits CaN, and FK506, which forms a complex with the FKBP12 protein.

Before the experiment, the cells were pre-incubated for 1 h with 20 μM CsA or 5 μM FK506. Then the Cl^-^ current was recorded between -120 and +160 mV from cells dialyzed using an internal solution containing 20 μM CsA or 5 μM FK506 to keep CaN blocked. Fig. 4A shows the TMEM16A current-voltage relationships obtained from wild-type HEK293 cells untreated (green) and treated with 20 μM CsA (blue) or 5 mM FK506 (red). Both CaN inhibitors decreased TMEM16A currents to a similar degree. However, neither inhibitor altered the current density in cells that did not express CaN (Fig. 4B). Likewise, TMEM16A expressed in HEK293 cells transiently transfected with the scrambled shRNA against CaN were sensitive to 20 mM CsA (Fig. 4C) but not when the channel was expressed in cells transfected with the shRNA against CaN (Fig. 4D). These data demonstrate that inhibiting CaN with CsA or silencing CaN expression with the shRNA diminish channel activity. Therefore, it is plausible that CaN binds to the TMEM16A-FKBP12 complex to up-regulate the function of the TMEM16A channel.

**FIGURE 4.**
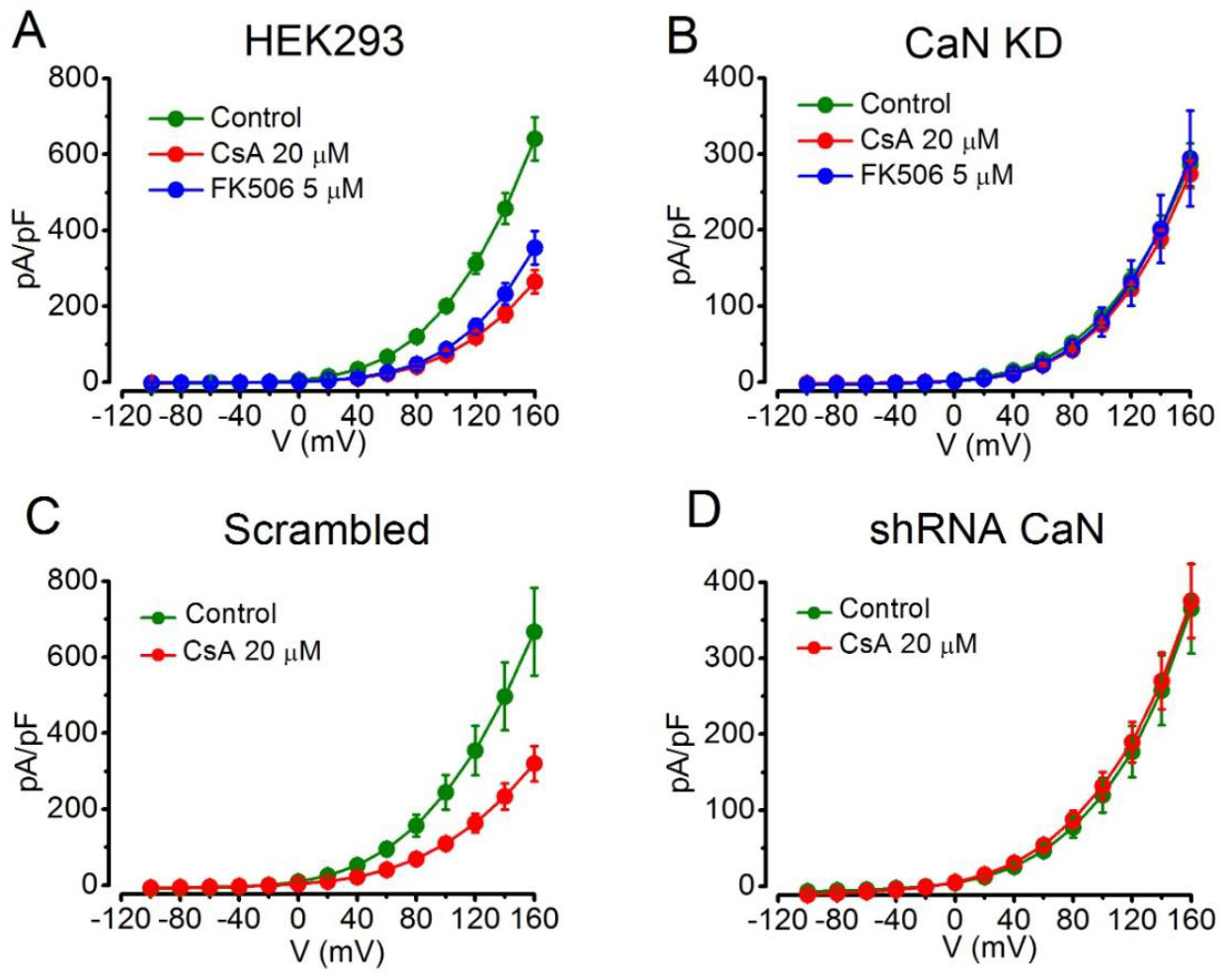
CaN regulates TMEM16A activity. **A)** Current-voltage relationships from TMEM16A expressed in wild-type HEK293 cells, **B)** HEK-AD293 cells stably transfected with shRNA CaN to knock-down the expression of CaN, **C)** wild-type HEK293 cells transiently transfected with scrambled shRNA CaN, and **D)** wild-type HEK293 cells transiently transfected with shRNA to decrease CaN expression. Cells untreated (green), exposed 1 h to 20 μM CsA (red), or exposed 1 h to 5 μM FK506 (blue).

## Discussion

In this work, we report that inhibiting CaN with CsA or FK506 diminishes the current magnitude through TMEM16A, indicating that CaN regulates the activity of TMEM16A. However, CaN is dispensable for the TMEM16A-FKBP12 interaction. Furthermore, we found that TMEM16A and FKBP12 colocalize. Data from a complementation fluorescent assay in living cells indicates that TMEM16A and FKBP12 come together, an idea supported by the *in-situ* interaction of recombinant FKBP12 with TMEM16A in isolated membranes. Based on this data, we propose that under physiological conditions, TMEM16A activity is supported by complexing with the heterodimer FKBP12-CaN.

In previous work, we speculated that the arrangement of the trimer was either TMEM16A-FKBP12-CaN or TMEM16A-CaN-FKBP12 (Sánchez-Solano et al., 2020). The present data show that TMEM16A-FKBP12 interaction is preserved in HEK293 cells devoid of CaN, indicating that the trimer is most likely to be TMEM16A-FKBP12-CaN. Colocalization and BiFC interaction data support the conclusion that TMEM16A interacts with FKBP12, even in basal Ca^2+^ conditions. Does FKBP12 have a role in controlling TMEM16A activity? The physiological role of FKBP12 in ion channel function is best illustrated by the effects documented on ryanodine receptors (RyR). In skeletal and cardiac muscles, FKBP12 helps maintain the function of RyR Ca^2+^ channels by directly interacting with the SPRY1 domain (Yuchi et al., 2015). A similar role may be fulfilled by FKBP12 when binding to the IP3 receptor Ca^2+^ channel (Cameron et al., 1995). Cardiomyopathy, cardiac hypertrophy, and heart failure arise when FKBP12 is absent (Wehrens et al., 2003), together with an increase in RyR single channel open probability that is accompanied by the development of sub-conductance states (Marx et al., 2000). Interestingly, the phosphorylation of RyR by PKA induces FKBP12 dissociation and increases the open probability and sub-conductance states. Additionally, RyR is phosphorylated by CaMKII; however, this phosphorylation event does not induce FKBP12 unbinding (Wehrens et al., 2004). Importantly, okadaic acid, a PP-1 and PP-2A phosphatases inhibitor, also increased the open probability of RyR, thus implying that the dephosphorylation of RyR is important for regulating the channel’s activity (Marx et al., 2000). It was shown that CaN binds to RyR via FKBP12 in a Ca^2+^-dependent manner, which enhanced Ca^2+^ release in a CsA-sensitive manner (Shin et al., 2002). On the other hand, the interaction of TMEM16A with FKBP12 and CaN shows some similarities to the RyR-FKBP12 interaction. For example, TMEM16A interacts with FKBP12, whereas CaN likely interacts in a Ca^2+^-dependent manner with TMEM16A via FKBP12, forming a heterotrimeric complex that is physiologically relevant (Sánchez-Solano et al., 2020). Because TMEM16A activity is downregulated by CaMKII phosphorylation (Ayon et al., 2019), whereas PP1, PP2A, and CaN phosphatases do the opposite (Greenwood et al., 2004a; Ayon et al., 2009), it is tempting to speculate that CaN plays a role in the phosphorylation/dephosphorylation equilibrium. In our hand, CsA and FK506 produced the same amount of inhibition of TMEM16A expressed in wild-type HEK293 cells, but their effects are blunted in HEK cells without CaN. Thus, we conclude that CsA and FK506 decrease TMEM16A activity by inhibiting the CaN activity and not by directly inhibiting TMEM16A. Whether the interaction of FKBP12 leads to some regulation of the channel’s functions remains to be determined; at this point, we think that the role of FKBP12 is to secure the binding of CaN to TMEM16A. By doing so, perhaps CaN is more efficient in maintaining TMEM16A in a dephosphorylated state, which enhances channel activity.

It is interesting to note that TMEM16A form a complex with IP3 in smooth muscle and dorsal root neurons (Akin et al., 2023; Jin et al., 2013), two FKBP12-sensitive proteins. Rapamycin, which also binds to FKBP12 without inhibiting CaN and disrupts FKBP12 interactions, inhibits DNA synthesis and smooth muscle cell growth (Marx et al., 1995). As has been extensively researched, TMEM16A plays an essential role in vascular smooth muscle, where it helps to control blood pressure (for recent reviews, see (Arreola et al., 2024, 2022)). Whether the inhibitory effect of rapamycin on smooth muscle is due to its ability to interrupt the interaction of FKBP12 with TMEM16A remains to be determined.

## Data Availability Statement

All the experimental data is included in this work.

## Acknowledgements

Research on TMEM16A channels is supported by the FORDECYT-PRONACES 1308052 2020 grant from CONAHCYT, Mexico.

Ana Elena López-Romero is supported by Graduate Student Fellowship 813742 from CONAHCYT, México. Miriam Huerta is supported by a Postdoctoral Fellowship from CONAHCYT, México. Rita Moran-Zendejas is supported by a Postdoctoral Fellowship from CONAHCYT, Mexico.

The authors thank Nancy Corral-Fernández, Carmen Hernández-Carballo, Mayra Delgado-Ramírez, Adán Guerrero, Alma Alva, and Eduardo Brito-Alarcón for their technical support, assistance with the BiFC plasmid designs and microscopy advice.

## Author contributions

María Luisa Guzmán-Hernández: Produced CaN KD cells, performed co-IP and BiFC assays, and wrote and reviewed the manuscript.

Miriam Huerta: Performed the co-localization analysis and wrote and reviewed the manuscript.

Ana E. López-Romero: Performed the patch clamp experiments, analyzed the data, and reviewed the manuscript.

Rita Morán-Zendejas: Performed the patch clamp experiments, analyzed the data, and reviewed the manuscript.

Abigail Betanzos: Performed the co-localization analysis and reviewed the manuscript.

Patricia Pérez-Cornejo: Analyzed the Co-IP and BiFC data, secured funds, and wrote and reviewed the manuscript.

Jorge Arreola: Analyzed all the data, secured funds, and wrote and reviewed the manuscript. The Authors declare no conflicts of interest in the content of this work.

## Notes

### Competing Interest Statement

The authors have declared no competing interest.

